# Control of bacterial population density with population feedback and molecular sequestration

**DOI:** 10.1101/225045

**Authors:** Reed D. McCardell, Shan Huang, Leopold N. Green, Richard M. Murray

## Abstract

Genetic engineering technology has become sophisticated enough to allow precise manipulation of bacterial genetic material. Engineering efforts with these technologies have created modified bacteria for various medical, industrial, and environmental purposes, but organisms designed for specific functions require improvements in stability, longevity, or efficiency of function. Most bacteria live in multispecies communities, whose composition may be closely linked to the effect the community has on the environment. Bacterial engineering efforts will benefit from building communities with regulated compositions, which will enable more stable and powerful community functions.

We present a design of a synthetic two member bacterial community capable of maintaining its composition at a defined ratio of [*cell type* 1]: [*cell type* 2]. We have constructed the genetic motif that will act in each cell in the two member community, containing an AHL-based negative feedback loop that activates ccdB toxin, which caps population density with increasing feedback strength. It also contains one of two ccdB sequestration modules, either the ccdA protein antitoxin, or an RNA device which prevents transcription and translation of ccdB mRNA, that rescues capped population density with induction. We compare absorbance and colony counting methods of estimating bacterial population density, finding that absorbance-based methods overestimate viable population density when ccdB toxin is used to control population density.

Prior modeling results show that two cell types containing this genetic circuit motif that reciprocally activate the other’s ccdB sequestration device will establish a steady state ratio of cell types. Experimental testing and tuning the full two member community will help us improve our modeling of multi-member bacterial communities, learn more about the strengths and weaknesses of our design for community composition control, and identify general principles of design of compositionally-regulated microbial communities.

## Introduction

Bacteria are major players in the function of the environments they inhabit, partially responsible for shaping immune responses in humans [1] and driving global geochemical cycles [2, 3]. Developments in DNA technologies, particularly DNA synthesis, molecular cloning, and sequencing, have created a major opportunity to genetically engineer bacteria to affect any of the many important phenomena they mediate, and potentially create new bacterial functions, Due to the speed of bacterial replication and enormous capacity of bacteria to colonize diverse niches, any genetic engineering efforts made with bacteria have the potential to be massively influential for the system in which they are deployed.

Early efforts to exploit the power of engineered bacteria in mammalian systems have led to the creation of engineered gut-commensal organisms capable of responding to chemical signals in the gut [4]. serving as long-term resident detectors of gut inflammation [5]. and protecting vaginal mucosa against HIV infection [6]. Other engineered bacteria have been designed to sequester atmospheric carbon dioxide and produce carbon-neutral energy sources [7, 8]. or seek and detoxify the herbicide atrazine [9].

These successful efforts are important steps forward for practical microbiological engineering, but still want for improvements in stability, longevity, or efficiency of function [10, 11]. Many organisms identified in nature are not culturable in the laboratory, precluding their genetic manipulation [12, 13]. Additionally, bacteria do not usually mediate their effects on their environments as homogenous populations; they are almost always found as communities of different interacting species, strains and subtypes that often function better together than any single one would alone [14, 15]. The specific species composition of a bacterial community is a partial determinant of the community’s function [16, 17, 18, 19, 20], though regulation of the composition of a community by environmental factors and gene regulation by community members likely work together to control community function [21, 22]. Synthetic biologists have made great strides in the engineering of genetic circuits for dynamic regulation of gene expression within single cells, but engineering control of bacterial community composition presents the special challenge of linking the activity of a genetic circuit that exists inside individual cells to the number of cells of each type in a population.

Population control using genetic circuits has been achieved in a few forms over the last decade. You et al. created a single strain community capable of limiting its own population density [23] and Balagaddé et al. created a two member community that displays the oscillatory behavior of a predator-prey ecological relationship [24].

Diffusible aeyl-homoserinelaetone (AHL) quorum sensing signals are frequently used to allow information transfer between members of an engineered community [25, 26] and link the behaviors of cells. In population control circuits, AHL signals usually activate a population regulating effector, typically a genetically encoded toxin [27, 28, 11, 24, 23]. That said, other regulatory devices exist, including regulated transport of cross-feeding metabolites [29] or reciprocal activation of antibiotic resistance [30].

Motivated by the results of studies demonstrating remarkably stable compositions of various microbial communities over time in diverse environments [31, 32], we present a design of a two member community of bacteria that maintains a specific ratio between the densities of the two members in coculture: [*cell type A*] = *a*[*cell type B*] where *a* represents the desired proportionality relationship between the densities of *A* and *B*, with which we hope to learn basic design principles for heterogeneous microbial communities, Een et al. recently published a detailed model and analysis of this system, demonstrating that the proposed community architecture implements a lag compensator that can maintain a steady-state population composition robust to changes in cell number and strain growth rates [33].

Here we report the experimental characterization of the genetic circuit motif that will act in each cell type in the full two member community, which balances feedback activation of cell death via ccdB toxin and inducible repression of cell death via ccdB inhibition.

## Results

### Population Circuit Design

Each cell type in the two member community contains a symmetric genetic circuit that, at steady state, balances cell death from cis-acting AHL feedback activation of ccdB toxin and cell death inhibition from trans-acting AHL activation of a toxin sequestration device (Fig 1), Simulations and analysis of this system demonstrate its ability to maintain a steady community composition robust to various perturbations [33]. Toxin “sequestration” is differentiated from toxin “expression” by the biochemical level of regulation. Our sequestration devices do not affect the activity of the promoter driving transcription of ccdB, but “sequester” either mRNA or ccdB protein, preventing transcription/translation of ccdB at the mRNA level [34]. or binding ccdB protein in a nonfunctional complex [35, 36], respectively.

**Figure 1:**
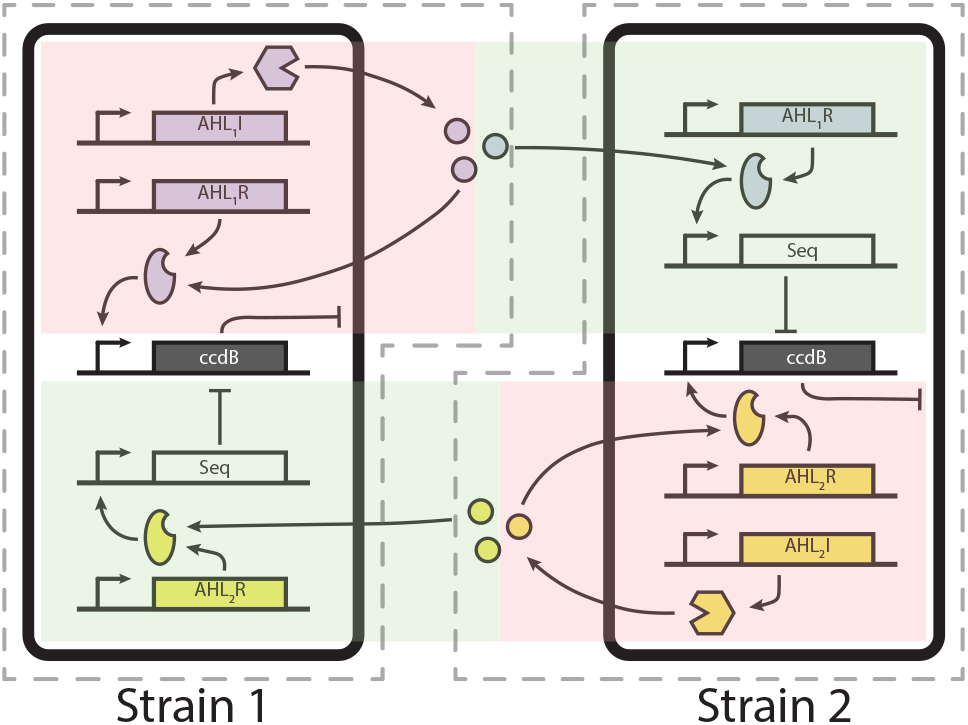
Schematic of two coll composition control circuit. Dashed outlines separate symmetric halves of the circuit that interact through two different AHL molecules. Components shaded red are involved in feedback activation of ccdB toxin and cell death. Components shaded green are involved in rescue from cell death by a sequestration device (Seq).

We explore 2 types of toxin sequestration devices, one being ccdB’s protein antitoxin ccdA, the other a transcription/translation-inhibiting RNA device (RNA-IN and RNA-OUT) [34]. The ccdA/B pair combines both cytotoxic action and the sequestration mechanism, preventing exploration of other toxins that do not have a protein antitoxin. The RNA-IN/OUT system separates sequestration and cytotoxicity modules. allowing future exploration of arbitrary toxins in place of ccdB.

Before creating the complete community from Fig 1, we considered the circuit motif in Fig 2 and simulated its ability to affect the density of a single cell type in monoculuture (Fig 2), The circuit has 2 modules: a negative feedback loop causing self-limitation of cell density through expression of ccdB toxin (population capping); and a toxin sequestration module, which can be activated by external inducer to allow growth to a higher density than allowed by the negative feedback arm (population rescue). The circuit can be conceptualized as 2 dials, a red dial that regulates the strength of AHL feedback on the cell and thus the steady state population density in the absence of toxin sequestration. Higher numbers on the dial correspond to lower steady state population densities. The green dial regulates the amount of toxin sequestration and thus, the amount to which a culture is allowed to increase its density above the steady state set by the negative feedback arm. Higher numbers correspond to higher steady state population densities.

**Figure 2:**
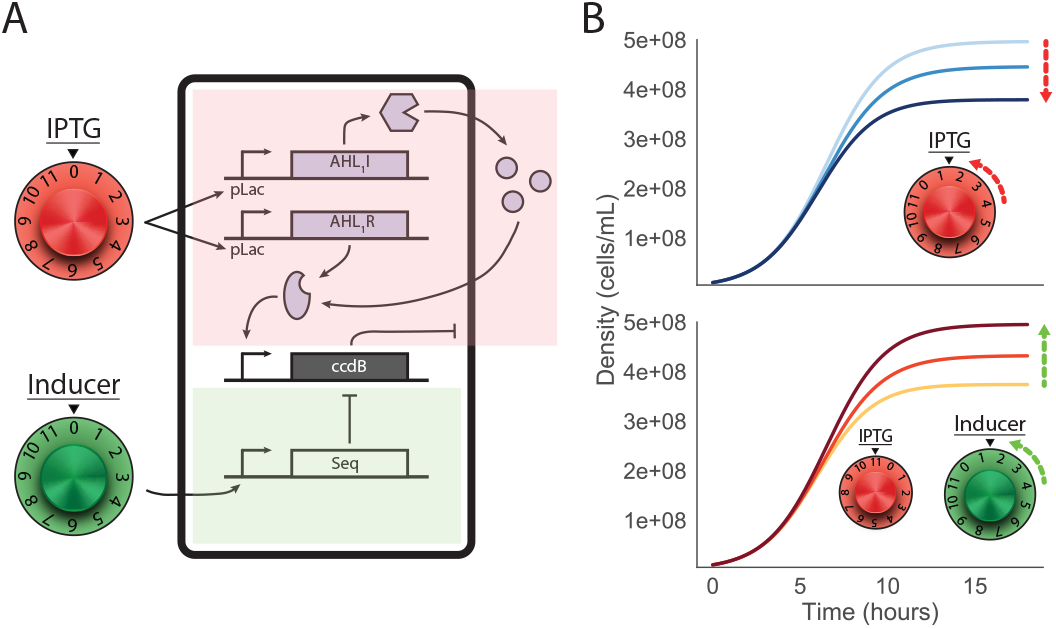
Schematic of single cell density control circuit. As in Fig 1. red shading indicates participation in feedback toxin activation, green indicates toxin sequestration. The red dial represents the level of induction of the indicated feedback components. Turning the red dial by adding IPTG increases expression of feedback components and the strength of feedback toxin activation. Similarly, turning the green dial by adding the appropriate inducer increases expression of the toxin sequestration device. See Materials and Methods “Modeling and Simulations” for modeling detail.

### Experimental Data

To verify the two functions of our genetic circuit motif, we transformed *E. coli* cells with the components in Fig 2 and incubated a dilute suspension of cells in varying concentrations of isopropyl *β*-D-l-thiogalactopvranoside (IPTG) for 18 hours, to induce expression of the AHL components involved in feedback population density limitation and observe the effect of population feedback on steady state population density. Induction of population capping by IPTG leads to reduction in steady state population density in circuits containing each ccdB sequestration module, as measured by absorbance at 600nm (Fig 3).

**Figure 3:**
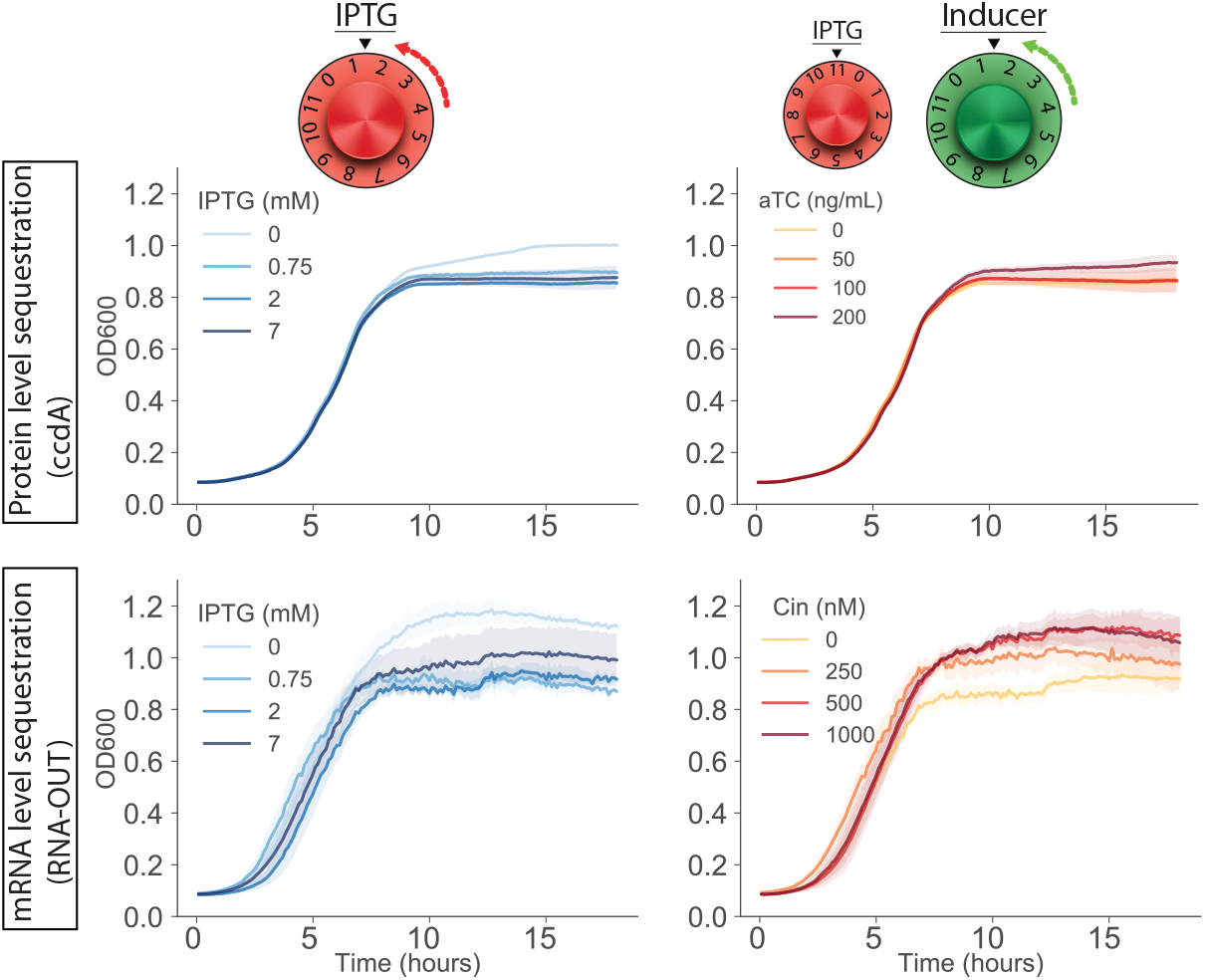
*E. colt* transformed with the circuit components in Fig 2 were induced with IPTG (bine curves) and the appropriate rescue inducer (red curves) to induce population capping and rescue, respectively. The top row contains ccdA protein as the ccdB sequestration module, inducible by aTC. The bottom row contains RNA-IN + ccdB as the ccdB niRNA and RNA-OUT as the ccdB sequestration module, inducible by Cin AHL. Each curve represents the mean of 3 replicates, shaded regions represent the standard deviation.

We confirmed the ability of each of our ccdB sequestration modules to rescue steady state population density above the feedback limited density by incubating cells in the maximum concentration of IPTG and varying concentrations of the appropriate inducer to express each sequestration module, A population strongly induced by IPTG and various amounts of secondary inducer establishes a higher steady state density than a population induced by IPTG alone (Fig 3).

The function of the circuit motif is affected by the ccdB sequestration device driving it. In the circuit variant with protein-level sequestration (ccdA, top row in Fig 3), growth of each population is smooth and consistent across replicates. In the variant employing RNA level sequestration (RNA-IN/OUT, bottom row in Fig 3), growth of each population is jerky and noisy across replicates. We suspect the level of ccdB sequestration is responsible for these effects; at lower mRNA copy numbers compared to protein copy numbers, stochastic sequestration may play a larger role in regulating ccdB activity, producing the observed noise and variability in cell growth.

The steady state values of OD600 absorbance at increasing levels of IPTG are only modestly lower than the uninduced steady state, though previously reported results with similar AHL feedback activation of ccdB demonstrate 10-fold reduction in cell counts [23]. These results imply that population negative feedback is too weak in our circuit motif and limit our ability to demonstrate many, significantly different density steady states with feedback limitation or sequestration rescue. We suspected that the cytotoxic mechanism of ccdB does not kill a cell in a way that reduces its contribution to OD600, rather, dead cells may continue to absorb strongly at 600nm [36, 37, 38]. We compared OD600 absorbance values to viable cell counts obtained with two methods of culture plating and counting of colony forming units (CFU). Viable cell counts in cultures strongly induced with IPTG revealed drastically lower population densities relative to uninduced cultures than are captured by OD600 absorbance. Population steady states produced during rescue from capping were similarly found to be lower than uncapped steady states when measured by CFU counting (Fig 4).

**Figure 4:**
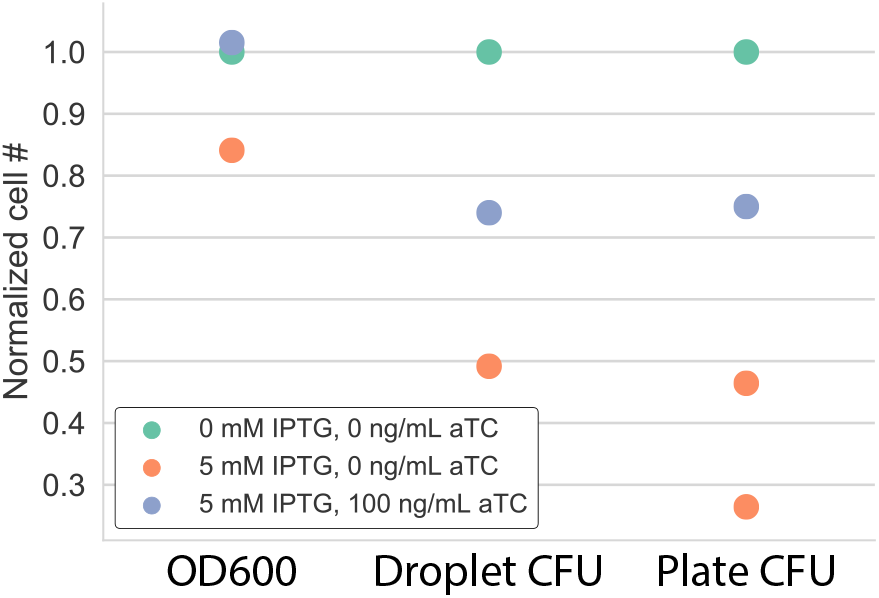
3 methods of cell density quantification were used to determine cell density after inducing population capping (**red**) and rescue (**blue**) in cells transformed with the circuit motif containing the protein-level toxin sequestration module. Values for each column are self-normalized to the absolute value of the **green** condition. Cell densities are those achieved after 18 hours of growth in the experimental conditions.

## Discussion

We demonstrate the ability of a genetic circuit motif employing negative feedback activation of ccdB toxin and inducible sequestration of ccdB to both negatively and positively regulate the steady state density of a bacterial culture. Two methods of ccdB sequestration that act on different biochemical levels were tested and shown to sequester and inhibit ccdB in the circuit context, through more precise experimentation is needed to compare their relative strengths of ccdB inhibition. While the circuit design using ccdB and ccdA is not modifiable to use different toxins except those with protein antitoxins, a circuit designed using RNA sequestration can be modified to function with arbitrary genes.

On its own, this circuit motif requires more testing in dynamic environments, that is, when the different inducers change concentration during cell growth. Additionally, it will be important to verify the claims that this circuit motif can reject perturbations in cell number and cell growth rate. We have established automated protocols to create arbitrary dynamics in inducer concentrations, perturbations to the number of cells in culture and perturbations in cell growth rate.

This circuit motif is designed to be linked between two cell types to maintain a stable ratio between the densities of each cell type. This system requires the use of two orthogonal AHL quorum sensing systems to allow the feedback capping and rescue modules in each cell to function without crosstalk. The circuit motifs tested in this study will be modified to include the appropriate receptors, synthases and promoters to create the community circuit presented in Fig 1.

More data is needed to precisely determine the amount to which OD600 overestimates viable cell counts, but it is clear that cell counting methods uncover a more realistic estimate of viable population density. We have added the droplet-based CFU counting method to our automated experimental protocols and hope to adjust our models of cell density to reflect the inaccuracy of OD600 measurements to allow maximal inference of circuit function from highly time-resolved OD600 data.

Testing and tuning the full two member community will help us learn more about the strengths and weaknesses of our design for community composition control and identify general principles of design of compositionally-regulated microbial communities.

## Materials and Methods

### *E. coli* cell strains

*E. coli* strain DH5αZ1 was used for the creation of the cell strain containing the circuit motif with protein-level ccdB sequestration; strain CY027 [39] was used to create the cell strain containing the motif with RNA-level ccdB sequestration. Both strains have genome integrations expressing the necessary activator/repressor transcription factors to allow regulated expression of circuit components: DH5αZ1 has genome integrated expression of LacI and TetR; C027 has genome integrated expression of both RhlR and CinR.

### Plasmids

The circuit motif with protein-level ccdB sequestration contains 3 plasmids: pLuxRI2, plw,:rCcdB3 (both from [23]), and pTetCcdA, pTetCcdA was constructed by GoldenGate assembly of the pTet promoter, the B0033m weak RBS, the *ccdB* gene taken from pOSIP_KO plasmid [40], and the B0015 terminator into a pSC101 backbone containing carbenicillin resistance.

The circuit motif with RNA-level ccdB sequestration contains 3 plasmids: pRNAINc-cdB, pRNAOUT and pRhll, pRNAINeedB was constructed by Gibson assembly of the pRhl promoter, the RNA-IN module [34]. the *ccdB* gene and the B0015 terminator into a p15a backbone containing chloramphenicol resistance. pRNAOUT was constricted by Gibson assembly of the pCin promoter and the RNA-OUT module into a ColE1 backbone containing kanamycin resistance, pRhll was constructed by Gibson assembly of the J23106 promoter, the B0034 strong RBS, *lacI* gene, and B0015 terminator; the pLac promoter, B0034 strong RBS, the *rhlI* gene and the B0015 terminator into a pSCl01 backbone containing carbenicillin resistance.

Unless otherwise noted, all parts used in cloning come from the CIDAR MoClo parts kit [41]. The CIDAR MoClo Parts Kit was a gift from Douglas Densmore (Addgene kity # 1000000059), The pOSIP plasmid kit used for clonetegration was a gift from Drew Endy and Keith Shearwin (Addgene kit # 1000000035).

### Cell Growth Experiments

Cells containing a circuit motif variant were grown from a frozen glycerol stock in TBK media (10g tryptone, 7g KCl per liter, 100mM MOPS buffer) overnight, then were diluted 200x into fresh TBK media with carbenicillin (100μg/mL), kanamycin (50μg/mL) and chlorampenicol (25μg/mL) and aliquoted in triplicate in 500uL into a square 96 well Matriplate (dot Scientific, MGB096-1-1-LG-L), The plate was incubated for 18 hours in a Biotek Synergy H2 incubator/plate reader at 37°C while OD600 measurements were taken every 7 minutes.

Inducers were added to the 96 well Matriplate before cell suspensions were aliquoted, A Labeyte Echo 525 Liquid Handler was used to aliquot inducers into each well of the plate before cell suspensions were added.

### Cell Density Quantification

OD600 measurements were taken every 7 minutes during the growth period. CFU measurements were taken once at the end of an 18 hour growth cycle. OD600 measurements were taken using a Biotek Synergy H2 incubator/plate reader. Colony forming units were counted using two methods:

Droplet CFU counting: Cell suspensions were diluted 25,000x into fresh TBK media and aliquoted into a Labcyte Echo 384 well source plate. 50μL drops of this suspension were transferred to a Nunc OmniTray (ThermoFisher: 140156) filled with LB agar containing the appropriate antibiotics. The OmniTray was incubated at 37¼C overnight, then colonies were counted. The fraction of droplets spotted on the plate that DID NOT grow colonies was fit to a Poisson distribution to determine λ, which yielded the mean cells/mL.

Plate CFU counting: Cell suspensions were diluted 1 * 10^6^x into fresh TBK media, then 10-50uL of this suspension was spread on LB agar petri dishes. These plates were incubated at 37°C then colonies were counted. The number of colonies grown was multiplied by the dilution factor to obtain cells/mL.

#### Modeling and Simulations

We constructed a general ODE model of the circuit motif under study. The model does not currently capture the biophysical details of the different sequestration devices and is meant to be a high level exploration of the sequestration concept. The curves in Fig 2 are simulated from this model, by varying *k_E_* (Fig 2B top) to simulate stronger AHL feedback and *k_T_* (Fig 2B bottom) to simulate stronger induction of the sequestration device.

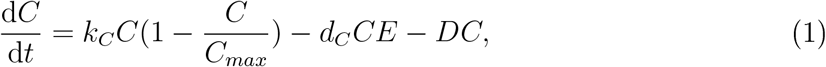

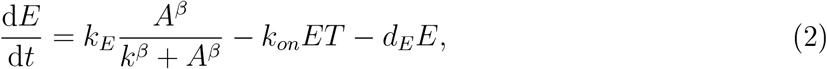

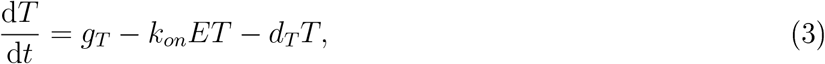

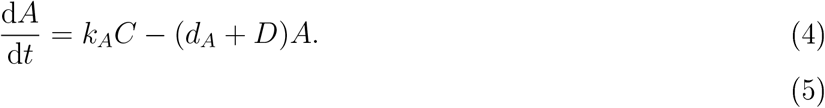

#### The variables in this model are as follows

C: :cell density 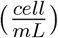
E: :CcdB concentration (*nM*)
T: :Sequestration device concentration (*nM*)
A: :AHL concentration (*nM*)

#### Parameters in the model

*k_c_*: :cell growth rate constant (0.01 – 0.04 *min*^−1^)
*C_max_*: :carrying capacity for cell growth (10^9^ *ml*^−1^)
*β*: :eooperativity of AHL effect (*β* = 2)
*d_c_*: :cell death rate contant by CcdB (2 × 10^−3^ *nM*^−1^ · *min*^−1^)
*k*: :concentration of AHL to half-maximally active promoter (10 *nM*)
*D*: :dilution rate (0.01 *min*^−1^)
*k_on_*: :binding rate of CcdB and sequestration device (0.05 *nM*^−1^ · *min*^−1^)
*g_T_, g_R_*: :basal production rate (0.01 – 0.1 *nM* · *min*^−1^)
*k_E_, k_T_*: :synthesis rate constant of CcdB, sequestration device (0.01 – 0.1 *nM* · *min*^−1^)
*k_A_*: :synthesis rate constant of AHL (5 × 10^−9^ *nM* · *ml* · *min*^−1^)
*d_A_*: :decay rate constant of AHL (0.01 *nM* · *min*^−1^)

Parameter estimates were found in a few literature sources [24, 23, 42]

In Fig 2, intial conditions are:

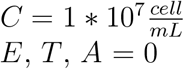

## Acknowledgments

The authors would like to thank Ania-Ariadna Baetica and Xinying Ren for their mathematical insights into the studied systems. The authors also thank Mark Prator for his assistance in performing the presented experiments. This project is sponsored by the Defense Advanced Research Projects Agency (Agreement HR0011-17-2-0008). The content of the information does not necessarily reflect the position or the policy of the Government, and no official endorsement should be inferred.

## References

[1] J. L. Round and S. K. Mazmanian, “The gut microbiome shapes intestinal immune responses during health and disease,” Nature Reviews. Immunology, vol. 9, pp. 313–323, may 2009.

[2] P. Flombaum, J. L. Gallegos, R. A. Gordillo, J. Rincon, L. L. Zabala, N. Jiao, D. M. Karl, W. K. W. Li, M. W. Lomas, D. Veneziano, C. S. Vera, J. A. Vrugt, and A. C. Martiny, “Present and future global distributions of the marine Cyanobacteria Proehloro-coccus and Syneehoeoeeus,” Proceedings of the National Academy of Sciences of the United States of America, vol. 110, pp. 9824–9829, jun 2013.

[3] W. E. Newton, “Biology of the nitrogen cycle,” ch. Physiology, Biochemistry, and Molecular Biology of Nitrogen Fixation, pp. 109–129, Amsterdam: Elsevier, 2007.

[4] M. Mimee, A. C. Tucker, C. A. Voigt, and T. K. Lu, “Programming a Human Commensal Bacterium, Bacteroides thetaiotaomicron, to Sense and Respond to Stimuli in the Murine Gut Microbiota,” Cell Systems, vol. 1, no, 1, pp. 62–71, 2015.

[5] D. T. Riglar, T. W. Giessen, M. Baym, S. J. Kerns, M. J. Niederhuber, R. T. Bronson, J. W. Kotula, G. K. Gerber, J. C. Way, and P. A. Silver, “Engineered bacteria can function in the mammalian gut long-term as live diagnostics of inflammation,” Nature Biotechnology, vol. 35, p. 653, may 2017.

[6] L. A. Lagenaur, B. E. Sanders-Beer, B. Brichacek, R. Pal, X. Liu, Y. Liu, R. Yu, D. Venzon, P. P. Lee, and D. H. Hamer, “Prevention of vaginal SHIV transmission in macaques by a live recombinant Lactobacillus,” Mucosal Immunology, vol. 4, p, 648, jul 2011.

[7] C. Liu, B. C. Colón, M. Ziesack, P. A. Silver, and D. G. Nocera, “Water splitting biosynthetic system with C02 reduction efficiencies exceeding photosynthesis,” Science, vol. 352, no. 6290, pp. 1210–1213, 2016.

[8] M. Kanno, A. L. Carroll, and S. Atsumi, “Global metabolic rewiring for improved C02 fixation and chemical production in cyanobacteria,” Nature Communications, vol. 8, p. 14724, mar 2017.

[9] J. Sinha, S. J. Reyes, and J. P. Gallivan, “Reprogramming bacteria to seek and destroy an herbicide,” Nature Chemical Biology, vol. 6, p. 464, may 2010.

[10] M. S. Donia, “A Toolbox for Mierobiome Engineering,” Cell Systems, vol. 1, pp. 21–23, nov 2017.

[11] F. K. Balagaddé, L. You, C. L. Hansen, F. H. Arnold, and S. R. Quake, “Long-term monitoring of bacteria undergoing programmed population control in a mieroehemo-stat,” Science, vol. 309, no. 5731, pp. 137–140, 2005.

[12] E. J. Stewart, “Growing uneulturable bacteria.,” Journal of bacteriology, vol. 194, pp. 4151–4160, aug 2012.

[13] A. W. Walker, S. H. Duncan, P. Louis, and H. J. Flint, “Phylogeny, culturing, and metagenomics of the human gut microbiota.” Trends in microbiology, vol. 22, pp. 267–274, may 2014.

[14] R. M. Stubbendieck, C. Vargas-Bautista, and P. D. Straight, “Bacterial Communities: Interactions to Scale,” Frontiers in Microbiology, vol. 7, p. 1234, aug 2016.

[15] H. C. Chung, O. O. Lee, Y.-L. Huang, S. Y. Mok, R. Kolter, and P.-Y. Qian, “Bacterial community succession and chemical profiles of subtidal biofilms in relation to larval settlement of the polychaete Hydroides elegans,” The ISME Journal, vol. 4, p. 817, jan 2010.

[16] J. A. Gilbert, R. A. Quinn, J. Debelius, Z. Z. Xu, J. Morton, N. Garg, J. K. Jansson, P. C. Dorrestein, and R. Knight, “Microbiome-wide association studies link dynamic microbial consortia to disease.” Nature, vol. 535, no. 7610, pp. 94–103, 2016.

[17] A. B. Shreiner, J. Y. Kao, and V. B. Young, “The gut mierobiome in health and in disease,” Curr Opin Gastroenterol, vol. 31, pp. 69–75, jan 2015.

[18] P. J. Turnbaugh, R. E. Ley, M. A. Mahowald, V. Magrini, E. R. Mardis, and J. I. Gordon, “An obesity-associated gut microbiome with increased capacity for energy harvest,” Nature, vol. 444, pp. 1027–1131, dec 2006.

[19] D. R. Nemergut, A. Shade, and C. Violle, “When, where and how does microbial community composition matter?,” Frontiers in Microbiology, vol. 5, p. 497, sep 2014.

[20] M. S. Strickland, C. Lauber, N. Fierer, and M. A. Bradford, “Testing the functional significance of microbial community composition,” Ecology, vol. 90, pp. 441–451, feb 2009.

[21] C. L. Dupont, J. P. McCrow, R. Valas, A. Moustafa, N. Walworth, U. Goodenough, R. Roth, S. L. Hogle, J. Bai, Z. I. Johnson, E. Mann, B. Palenik, K. A. Barbeau, J. Craig Venter, and A. E. Allen, “Genomes and gene expression across light and productivity gradients in eastern subtropical Pacific microbial communities,” The ISME Journal, vol. 9, pp. 1076–1092, may 2015.

[22] A. Fernandez, S. Huang, S. Seston, J. Xing, E. Hickey, C. Criddle, and J. Tiedje, “How Stable Is Stable? Function versus Community Composition,” Applied and Environmental Microbiology, vol. 65, no. 8, pp. 3370–3697, 1999.

[23] L. You, E. S. Cox, E. Weiss, and F. H. Arnold, “Programmed population control by cell-cell communication and regulated killing,” Nature, vol. 428, no. 6985, pp. 868–871, 2004.

[24] F. K. Balagaddé, H. Song, J. Ozaki, C. H. Collins, M, Barnet, F. H. Arnold, S. E. Quake, and L. You, “A synthetic Escherichia coli predator-prey ecosystem.,” Molecular Systems Biology, vol. 4, no. 187, p. 187, 2008.

[25] S. E. Scott and J. Hasty, “Quorum Sensing Communication Modules for Microbial Consortia,” ACS Synthetic Biology, vol. 5, no. 9, p. aessynbio,5b00286, 2016.

[26] M. B. Miller and B. L. Bassler, “Quorum Sensing in Bacteria,” Annual Review of Microbiology, 2001.

[27] S. E. Scott, M. O. Din, P. Bittihn, L. Xiong, L. S. Tsimring, and J. Hasty, “A stabilized microbial ecosystem of self-limiting bacteria using synthetic quorum-regulated lvsis,” Nature Microbiology, vol. 2, p. 17083, jun 2017.

[28] P. Marguet, Y. Tanouchi, E. Spitz, C. Smith, and L. You, “Oscillations by Minimal Bacterial Suicide Circuits Eeveal Hidden Facets of Host-Circuit Physiology,” PLoS ONE, vol. 5, p. ell909, jul 2010.

[29] A. Kerner, J. Park, A. Williams, and X. N. Lin, “A programmable escherichia coli consortium via tunable symbiosis,” PLoS ONE, vol. 7, no. 3, pp. 1–10, 2012.

[30] B. Hu, J. Du, E. yang Zou, and Y. J. Yuan, “An environment-sensitive synthetic microbial ecosystem,” PLoS ONE, vol. 5, no. 5, pp. 1–9, 2010.

[31] C. Huttenhower and Human Microbiome Project Consortium, “Structure, function and diversity of the healthy human microbiome.,” Nature, vol. 486, no. 7402, pp. 207–14, 2012.

[32] M. Musilova, M. Tranter, S. A. Bennett, J. Wadham, and A. M. Anesio, “Stable microbial community composition on the Greenland Ice Sheet,” Frontiers in Microbiology, vol. 6, no. MAE, pp. 1–10, 2015.

[33] X. Een, A.-A. Baetica, A. Swaminathan, and E. M. Murray, “Population regulation in microbial consortia using dual feedback control,” To appear, IEEE Conference on Decision and Control. bioRxiv, 2017. https://doi.org/10.1101/120253.

[34] C. C. Liu, L. Qi, J. B. Lucks, T. H. Segall-Shapiro, D. Wang, V. K. Mutalik, and A. P. Arkin, “An adaptor from translational to transcriptional control enables predictable assembly of complex regulation,” Nature Methods, vol. 9, no. 11, pp. 1088–1094, 2012.

[35] E. M. Bahassi, M. H. O’Dea, N. Allali, J. Messens, M. Gellert, and M. Couturier, “Interactions of CcdB with DNA: Gyrase in activation of GyrA, poisoning of the gvrase-DNA complex, and the antidote action of CdcA,” Journal of Biological Chemistry, vol. 274, no. 16, pp. 10936–10944, 1999.

[36] N. De Jonge, A. Garcia-Pino, L. Buts, S. Haesaerts, D. Charlier, K. Zangger, L. Wyns, H. De Greve, and E. Loris, “Rejuvenation of CedB-Poisoned Gvrase by an Intrinsically Disordered Protein Domain,” Molecular Cell, vol. 35, no. 2, pp. 154–163, 2009.

[37] S. E. Critchlow, M. H. O’Dea, A. J. Howells, M. Couturier, M. Gellert, and A. Maxwell, “The interaction of the F plasmid killer protein, CcdB, with DNA gyrase: induction of DNA cleavage and blocking of transeriptionllJ. Karn,” Journal of Molecular Biology, vol. 273, no. 4, pp. 826–839, 1997.

[38] M.-H. Dao-Thi, L. Van Melderen, E. De Genst, H. Λlif. L. Buts, L. Wvns, and E. Loris, “Molecular Basis of Gyrase Poisoning by the Addiction Toxin CcdB,” Journal of Molecular Biology, vol. 348, no. 5, pp. 1091–1102, 2005.

[39] Y. Chen, J. K. Kim, A. J. Hirning, R. Matthew, K. Josi, and M. R. Bennett, “Emergent genetic oscillations in a synthetic microbial Consortium,” Science, vol. 349, no. 6251, pp. 986–989, 2016.

[40] F. St-Pierre, L. Cui, D. G. Priest, D. Endy, I. B. Dodd, and K. E. Shearwin, “One-step cloning and chromosomal integration of DNA,” ACS Synthetic Biology, vol. 2, no. 9, pp. 537–541, 2013.

[41] S. V. Iverson, T. L. Haddock, J. Beal, and D. M. Densmore, “CIDAR MoClo: Improved MoClo Assembly Standard and New E. coli Part Library Enable Rapid Combinatorial Design for Synthetic and Traditional Biology,” ACS Synthetic Biology, vol. 5, no. 1, pp. 99–103, 2016.

[42] R. Smith, C. Tan, J. K. Srimani, A. Pai, K. A. Riccione, H. Song, and L. You, “Programmed Allee effect in bacteria causes a tradeoff between population spread and survival,” Proceedings of the National Academy of Sciences of the United States of America, vol. Ill, pp. 1969–1974, feb 2014.

